# CD3, CD28, TCRαβ expression and IL-2 production in a spontaneous glycosylphosphatidylinositol-deficient Jurkat T cell line

**DOI:** 10.64898/2026.07.22.740193

**Authors:** William S Glass, Cindy L Zuleger, Yujia Cai, Michael A Newton, Mark R Albertini

## Abstract

Glycosylphosphatidylinositol (GPI) anchors are involved in the organization of membrane microdomains that support T cell receptor (TCR) signaling. However, their role in regulating expression of TCR-related proteins and downstream functional output remains unclear. This study aimed to characterize the effects of GPI-deficiency on TCR, cluster of differentiation 3 (CD3), and CD28 expression as well as interleukin-2 (IL-2) production using a GPI-deficient Jurkat T cell line (S12). Flow cytometry confirmed the complete loss of GPI anchors and GPI-anchored proteins (GPI-APs) in the S12 cell line. Compared to GPI-producing parental Jurkat, S12 had significantly higher expression of CD3 and TCRαβ while CD28 had similar expression. IL-2 production by S12 was assessed following stimulation with anti-CD3/anti-CD28 beads and following stimulation with phorbol 12-myristate 13-acetate (PMA) and ionomycin. Neither S12 nor parental Jurkat produced detectable IL-2 in response to anti-CD3/anti-CD28 bead-mediated stimulation. Both parental Jurkat and S12 produced IL-2 following PMA/ionomycin-mediated stimulation. No significant difference in IL-2 production was observed between S12 and parental Jurkat following PMA/ionomycin-mediated stimulation. These findings demonstrate that GPI-deficiency influences surface receptor expression but does not significantly impair downstream IL-2 production under PMA/ionomycin stimulation. This finding suggests that GPI anchors and GPI-APs contribute to proximal signaling organization but are not required for cytokine production when downstream pathways are directly activated.

## Introduction

Classical activation of naïve T cells requires T cell receptor (TCR) engagement with an antigenic peptide – major histocompatibility complex (MHC) molecule complex on an antigen presenting cell (1). Co-receptor-associated lymphocyte-specific protein tyrosine kinase (Lck) phosphorylates the immunoreceptor tyrosine-based activation motifs (ITAMs) on the cytosolic tails of cluster of differentiation 3 (CD3) (2). This phosphorylation leads to the docking and activation of zeta-chain-associated protein kinase 70 (ZAP70) (3). Activated ZAP70 phosphorylates linker for activation of T cells (LAT) and SH2 domain-containing leukocyte-specific phosphoprotein of 76 kDa (SLP-76), promoting the assembly of the signalosome that nucleates downstream signaling pathways (3). This signalosome recruits phospholipase Cγ 1 (PLCγ1), leading to inositol 1,4,5-trisphosphate (IP3)-dependent calcium mobilization and activation of calcineurin and nuclear factor of activated T cells (NFAT) while diacylglycerol (DAG)-dependent protein kinase Cθ (PKCθ) signaling activates nuclear factor κ-light-chain-enhancer of activated B cells (NF-κB) (1). Together, these activated transcription factors promote interleukin-2 (IL-2) production and T cell proliferation (1).

Complete T cell activation requires CD28 binding to CD80/CD86 on the antigen presenting cell in addition to TCR engagement (4). CD28 signaling enhances TCR signaling by amplifying phosphorylation-dependent assembly of signaling complexes at the immune synapse (4). CD28 engagement promotes phosphoinositide 3-kinase (PI3K) recruitment and, subsequently, phosphatidylinositol trisphosphate (PIP3) production at the immunological synapse (5). This engagement aids in recruitment of downstream activators including PLCγ1 and PKCθ, leading to activation of transcription factors like NF-κB (4). Overall, CD28 co-engagement enhances IL-2 transcription, metabolic fitness, and clonal proliferation (4).

Efficient T cell activation relies on the coordinated integration of TCR and CD28 signals. Due to this coordinated integration, the spatial organization of these receptors within membrane microdomains is critical for efficient signal propagation. Lipid rafts are dynamic microdomains within the plasma membrane that are enriched in glycosphingolipids, sphingomyelin, and cholesterol (6). This enrichment creates an environment that attracts certain proteins while excluding other proteins (6). Upon TCR ligation, these lipid rafts quickly coalesce and aid in the concentration of proximal signaling molecules like Lck, CD3, and LAT (7). Raft-associated recruitment of LAT is particularly important for the assembly of the LAT, SLP-76 signalosome that aids in activation of downstream activators previously mentioned (8). In addition to organizing proximal TCR signaling, these microdomains contribute to immune synapse stability by coordinating TCR and CD28 complexes, ultimately ensuring robust IL-2 production and proliferation (9).

Because membrane rafts depend on molecules that partition into and stabilize ordered lipid domains, it is suggested that GPI anchors and GPI-APs play an important role in T cell signaling. GPI-APs are proteins bound to the cellular membrane by GPI anchors and uniquely positioned to promote membrane compartmentalization through their organization in submicron domains (Fig 1) (10). These GPI-enriched domains are thought to promote nanoscale membrane organization and can contribute to the stability of ordered membrane regions (10, 11). Based on this stabilization, it is proposed that disruption of GPI anchor biosynthesis has the potential to alter the membrane compartmentalization involved in T cell activation.

**Fig 1.**
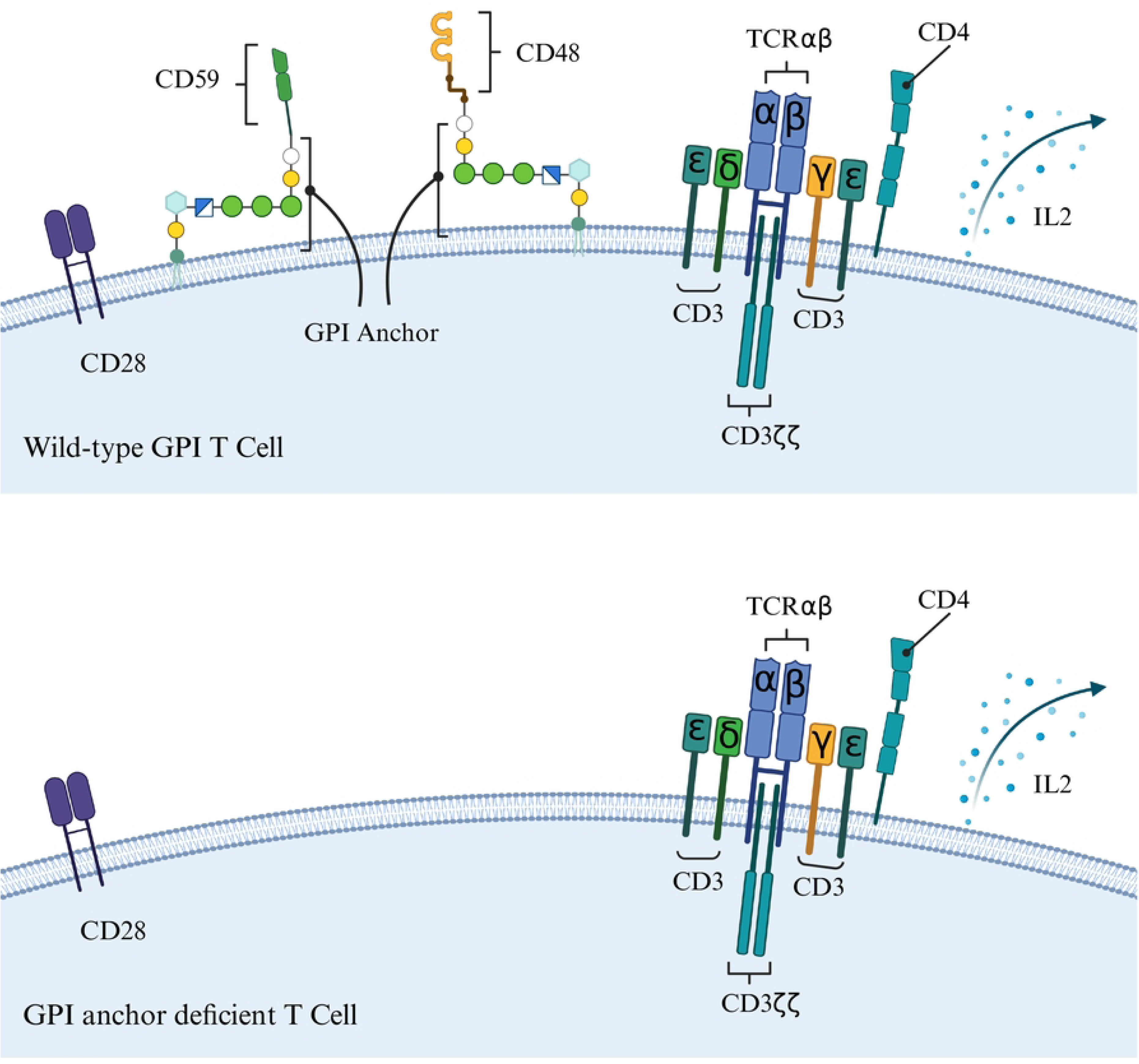
Wild-type and GPI deficient T cell phenotype. Wild-type T cells express GPI anchors that link certain proteins, e.g., CD48 and CD59, to the cellular membrane while GPI deficient T cells lack them.

Prior studies support a functional role for GPI anchors and GPI-APs in T cell activation and its outputs. Early work demonstrated that engagement of the GPI-AP CD59 on human T cells induced IL-2 production, but only in the presence of an intact CD3/TCR complex (12). This IL-2 production indicates there is a signaling pathway between GPI-APs and the TCR during canonical TCR signaling. Additional studies on GPI-deficient T cells derived from patients with paroxysmal nocturnal hemoglobinuria (PNH) showed that proliferative responses to anti-CD3 stimulation are significantly reduced despite these T cells having preserved surface expression of CD3ε and comparable levels of CD4 (13). PNH is caused by somatic mutation in the X-linked phosphatidylinositol glycan class A (*PIGA*) gene, and the PIGA protein is critical early in the biosynthetic pathway of GPI anchors (14). However, it remains unclear how loss of GPI anchors and loss of GPI-APs influence surface expression of TCRαβ, CD3, and CD28. It is also unclear how these changes relate to IL-2 production. To address this question, a spontaneous GPI-deficient Jurkat T cell line isolated in our laboratory and referred to as “S12” was examined for surface expression of TCRαβ, CD3, and CD28 and assessed for IL-2 production following T cell stimulation.

## Materials and methods

### Cell culture

The Jurkat, clone E6-1, human T cell leukemia cell line (RRID:CVCL_0367) was authenticated by the University of Wisconsin Translational Research Initiatives in Pathology (TRIP) Laboratory (Table 1) and cultured in RPMI-1640 supplemented with 10% fetal bovine serum (FBS) (BenchMark™, GeminiBio), 25 mM HEPES, 2 mM L-glutamine, 1 mM sodium pyruvate, and 1X non-essential amino acids (all from Corning Inc.) and 50 μM 2-mercaptoethanol (Sigma-Aldrich) – hereto, referred to as R-10%. Jurkat was cultured in Falcon 75 cm^2^ tissue culture flasks (Corning) and maintained at 37°C in a humidified 5% CO_2_ atmosphere. The cell line S12, a GPI anchor-deficient Jurkat subpopulation generated by single-cell flow sorting (described below), was cultured as described above for parental Jurkat.

**Table 1.** STR Analysis.

| **Table 1. STR Analysis** |  |  |
| --- | --- | --- |
| **STR Locus** | **Expected STRs** | **Observed STRs** |
| FGA | 20,21 | 20,21 |
| TPOX | 8,10 | 8,10 |
| D8S1179 | 13,14 | 13,14 |
| vWA | 18,18 | 18,18 |
| Amelogenin | X,Y | X,Y |
| Penta_D | 11,13 | 11,13 |
| CSF1PO | 11,12 | 10,11,12 |
| D16S539 | 11,11 | 11,11 |
| D7S820 | 8,12 | 8,12 |
| D13S317 | 8,12 | 8,12 |
| D5S818 | 9,9 | 9,9 |
| Penta_E | 10,12 | 10,12 |
| D18S51 | 13,21 | 13,21 |
| D21S11 | 31.2,33.2 | 31.2,32.2,33.2 |
| TH01 | 6,9.3 | 6,9.3 |
| D3S1358 | 15,17 | 15,17 |
| The Jurkat cell line sample used in this study matched the STR profile of the human cell line Jurkat (CVCL_0367) with 30/32 allelic match (93.8% match); thus, this sample is called a match to Jurat, Clone E6-1 (ATCC® TIB-152™).<br>STR, short tandem repeat |  |  |

### Human subjects and peripheral blood samples

The University of Wisconsin-Madison Health Sciences Institutional Review Board (IRB) approved this study (Protocol #2017-0409) on 08 February 2013. At the time of approval, the study ID number was 2012-0684. This protocol was given a new ID number 2017-0409 after continuing review. Participants were available for this study starting from the date of initial IRB approval (08 February 2013), and recruitment for this program of research is ongoing. Written informed consent was obtained and cells from a single participant were used in this study. Peripheral blood mononuclear cells (PBMCs) obtained by standard laboratory techniques were cryopreserved in medium containing 90% fetal calf serum (FCS) (GeminiBio) and 10% dimethylsulfoxide (Sigma-Aldrich).

### Detection of GPI-anchor and GPI-anchored proteins using flow cytometry

Cells were washed twice with PBS followed by incubation with Ghost Violet 510 (GV510) viability dye (Tonbo Biosciences) at 1:3,200 dilution in PBS for 15 minutes at 25°C. Cells were washed twice with wash buffer (2% FBS in PBS) by centrifugation at 300 x g for 5 minutes. Cells were then stained with a mixture of the recombinant protein FLAER-iFluor488 (1:320 dilution, Caprico Biotechnologies, catalog #310514), 1:320 dilution monoclonal mouse)-human CD48-Phycoerythrin (PE) (BioLegend, clone BJ40, RRID:AB_2075176), and 1:160 dilution monoclonal mouse anti-human CD59 Super Bright 600 (SB600) (Invitrogen, clone OV9A2, RRID:AB_2735008) diluted in wash buffer for 30 minutes at 4°C. FLAER is a recombinant protein that binds directly to the GPI anchor itself and is used to identify cells with disrupted GPI-AP expression. Cells were washed twice as above, fixed with BD Cytofix (BD Biosciences) for 20 minutes at 4°C, washed once as above, resuspended in wash buffer and stored at 4°C until acquisition. Fluorescent Minus One (FMO) controls consisting of cells stained with all reagents except either FLAER, anti-CD48, or anti-CD59 were used to inform gate placement. Data were acquired on a 5-laser full-spectrum Cytek Aurora (Cytek Biosciences) using Cytek Assay Settings in SpectroFlo software and single-stained Jurkat were used as spectral reference controls for unmixing. FlowJo v10 software (BD Biosciences) was used for data analysis.

### Single-cell flow sorting of spontaneous, GPI anchor-deficient Jurkat cells

Jurkat were stained as detailed in the prior section with the exception of fixation. Following the last wash, cells were resuspended in sterile wash buffer and kept on ice until sorted. Single live cells with low to nil FLAER, CD48-PE, and CD59 SB600 fluorescence were sorted into wells of a 96-well round-bottom plate filled with R-10% media using a MA900 Multi-Application Cell Sorter (Sony, Tokyo, Japan). Wells were checked for growth using an inverted microscope and actively growing clones were expanded and cryopreserved. GPI anchor-deficient phenotype was confirmed using flow cytometry as described in the previous section. The clone ‘S12’ was selected for further testing.

### CD3, CD28, & TCRαβ staining and flow cytometry

All Jurkat and S12 samples were assayed in triplicate. Jurkat and S12 were washed and stained with GV510 as previously described. Cells were then stained with a mixture of 1:400 anti-human CD3-Alexa Fluor 700 (BioLegend, clone UCHT1, RRID:AB_493740), 1:25 anti-human CD28-Brilliant Ultra Violet 395 (BUV395) (ThermoFisher, clone 28.2, RRID:AB_2920947), and 1:100 anti-human TCR*αβ*-allophycocyanin (APC) (ThermoFisher, clone IP26, RRID:AB_10597896) diluted in wash buffer for 30 minutes at 4°C. Cells were washed and fixed as described previously and stored at 4°C until acquisition. Fluorescent Minus One (FMO) controls consisting of cells stained with all reagents except either anti-CD3, anti-CD28, or anti-TCRαβ were used to inform gate placement. Data were acquired and analyzed on a 5-laser full-spectrum Cytek Aurora (Cytek Biosciences) using Cytek Assay Settings in SpectroFlo software and single-stained parental Jurkat were used as spectral reference controls for unmixing. FlowJo v10 software (BD Biosciences) was used for data analysis. The median fluorescence intensities (MFIs) of each marker in either Jurkat or S12 were exported from FlowJo.

### RNA Isolation

Total RNA was isolated from the PBS-washed pellets of parental Jurkat and S12 cell lines using the RNeasy Mini Kit and QIAShredders following manufacturer’s directions (Qiagen) without the optional DNase-treatment. A NanoDrop spectrophotometer (ThermoFisher) quantified RNA and DNA as well as assessed the purity of the material. The material was used for reverse transcription PCR (RT-PCR) of *PIGA* and as a source of genomic DNA (gDNA) for PCR of *PIGA* and cell line authentication.

### Primer design

Primers were designed using NCBI Primer-BLAST and features available on Integrated DNA Technologies (IDT) website. Primers for genomic (g) PCR were based on the NCBI Reference Sequence (NG_009786.1) for human *PIGA* gDNA. Primers for RT-PCR were based on the NCBI Reference Sequence (NM_002641.4) for human *PIGA* mRNA. All primers were purchased from IDT and resuspended in 1X TE buffer (Ambion) at 200 µM (stock). Working concentrations of 20 µM were made by a 1:10 dilution in ddH2O (Ambion). Aliquots of primers were stored at -20°C. All primers used are shown in Supplementary Table S1.

### PIGA genomic polymerase chain reaction (PCR)

The RNA extracted from parental Jurkat and S12 was not treated with the optional DNase and therefore this RNA was used as a source of gDNA. A 20 μl reaction consisted of 2 μl 10X PCR buffer (Qiagen), 0.4 μl 10 mM (each) dNTP (IDT), 0.8 μl 10 μM forward primer (IDT), 0.8 μl 10 μM reverse primer (IDT), 14.9 μL RNase-free H2O, 0.1 μl HotStarTaq Polymerase (Qiagen), plus 1 μl of Jurkat RNA/gDNA, S12 RNA/gDNA, whole human male gDNA (positive control) (Promega), or 1 μl RNase-free H_2_O (negative control). Each reaction received an equivalent amount (50 ng) of gDNA. Primers are listed in Supplementary Table S1. PCR cycles were 95°C (15 minutes (min)) then 35 cycles of: 94°C (30 seconds (sec)), 58°C (1 min), 72°C (90 sec), followed by 72°C (10 min) and a 4°C hold. Custal Omega was used for Multiple Sequence Alignment (MSA) (15).

### PIGA RT-PCR for middle of exon 2 through exon 6

*PIGA* has several alternate splice products in the very large exon 2. The smaller splice variants are “favored” in PCR and become the dominant products making analysis challenging. Therefore, RT-PCR was performed on the 3’ end of exon 2 through exon 6, bypassing the splice site. SuperScript III OneStep RT-PCR kit (Invitrogen) was used following manufacturer’s directions to first convert mRNA to cDNA. A 20 μl reaction consisted of 10 μl 2x Reaction Mix (Invitrogen), 0.4 μl 10 μM MIDF forward primer (IDT), 0.4 μl 10 μM Ex6R reverse primer (IDT), 7.4 μL RNase-free H2O, 0.8 μl SuperScript III RT/Platinum Taq Mix (Invitrogen), plus 1 μl of Jurkat RNA/gDNA, S12 RNA/gDNA, in-house total human RNA (positive control), or 1 μl RNase-free H_2_O (negative control). Each reaction received an equivalent amount (50 ng) of total RNA. Primers are listed in Supplementary Table S1. Cycling parameters were 55°C (45 min) and 94°C (2 min) then 35 cycles of: 94°C (15 sec), 58°C (30 sec), 68°C (105 sec), followed by 68°C (5 min) and a 4°C hold.

### Gel electrophoresis

PCR products were run on a 1.2% UltraPure agarose gel (Invitrogen) cast with 0.5 μg/ml ethidium bromide (BioRad) in 1x Tris-Borate-EDTA buffer (TBE) alongside either pGEM DNA Marker or 100 bp DNA Ladder. Images were captured using either a UV-based (FotoDyne, Inc) or green LED-based (Invitrogen) transilluminator system. Buffer and molecular weight markers from Promega.

### DNA sequencing

PCR products were sequenced using long-read amplicon sequencing on Oxford Nanopore’s PromethION platform at the University of Wisconsin-Madison Biotechnology Center DNA Sequencing Core Facility (RRID:SCR_017759). NCBI Reference Sequences NG_009786.1 and NM_002641.3 were used as the *PIGA* gDNA and mRNA reference sequence, respectively.

### T cell stimulation

Cryopreserved human PBMCs were thawed and resuspended at 1 x 10^6^ cells/ml in R10%. Parental Jurkat and S12 were resuspended at 1 x 10^6^ cells/ml in R-10%. Cell suspensions (0.5 ml per well) were added to a 24-well tissue culture plate (Costar #3524, Corning). Cells were unstimulated or stimulated with either 5 x 10^5^ Human T-Activator CD3/CD28 beads (1:1 cell:bead ratio) (ThermoFisher) or 25 ng/mL phorbol 12-myristate 13-acetate (PMA) + 1 μg/ml ionomycin (Sigma-Aldrich) in a total volume of 1 mL R-10%. Plates were incubated ∼20-22 hours at 37°C/5% CO_2_. Supernatants were collected, particulates removed by centrifugation, aliquoted and stored at -20°C. Three independent replicates of the stimulation were performed.

### IL-2 enzyme-linked immunosorbent assay (ELISA)

Tissue culture supernatants from the stimulation assays were thawed and assayed for IL-2 secretion using a Human IL-2 Quantikine ELISA kit following manufacturer’s directions (R&D Systems). Supernatants from PMA/Ionomycin stimulation were diluted 1:3 in buffer provided in the kit before assaying. Each supernatant, standard, and blank were assayed in triplicate. The optical density of each well was determined using a SpectraMax M3 Micro-Plate Reader (Molecular Devices) set to 450 nm with wavelength correction set to 540 nm. SoftMax Pro 7 software (Molecular Devices) was used to generate a four-parameter logistic (4-PL) standard curve fit and calculate the IL-2 concentration of each well after background subtraction of the assay blanks.

### Statistical analysis

Statistical analyses were performed in R version 4.5.3. For comparison of flow cytometric MFI between Parental Jurkat and S12, a fixed-effects linear model was fit with cell line as the factor of interest and run as a blocking factor. For IL-2 ELISA data, a mixed-effects two-factor ANOVA (cell line × stimulation) was fit with experiment as a fixed blocking factor and a random effect at the supernatant level to account for the clustering of triplicate wells from each preparation. Two response transformations were used, square-root (with negative values set to zero) and inverse hyperbolic sine (asinh). P-values < 0.05 were considered statistically significant. Plots were generated using R software version 4.5.3 with graphical packages (16, 17).

## Results

### Parental Jurkat matches the STR report of Jurkat, clone E6-1

STR analysis confirmed the parental Jurkat closely matched the American Type Culture Collection (ATCC) Jurkat, Clone E6-1 reference profile (Table 1). A 30/32 allelic match (93.8%) was observed, exceeding the established threshold for cell line authentication (18). Our parental Jurkat is a match to the Jurkat, Clone E6-1 (ATCC® TIB-152TM) cell line with the exception of the D21S11, and CSF1PO loci.

### S12 Jurkat cell line with spontaneous loss of GPI-anchored proteins

We sorted single, live cells from the parental Jurkat cell line with spontaneous loss of GPI-anchored proteins (see Fig 2A for gating strategy). Several clones were expanded *in vitro*. The loss of protein expression was confirmed in clone S12 by flow cytometry. GPI anchor expression was lower in S12 relative to the parental Jurkat (Figs 2B and 2C). S12 showed no detectable binding of FLAER, indicating GPI anchor loss, and lacked expression of the GPI-APs CD48 and CD59. In contrast, Jurkat showed robust expression of these markers.

**Fig 2.**
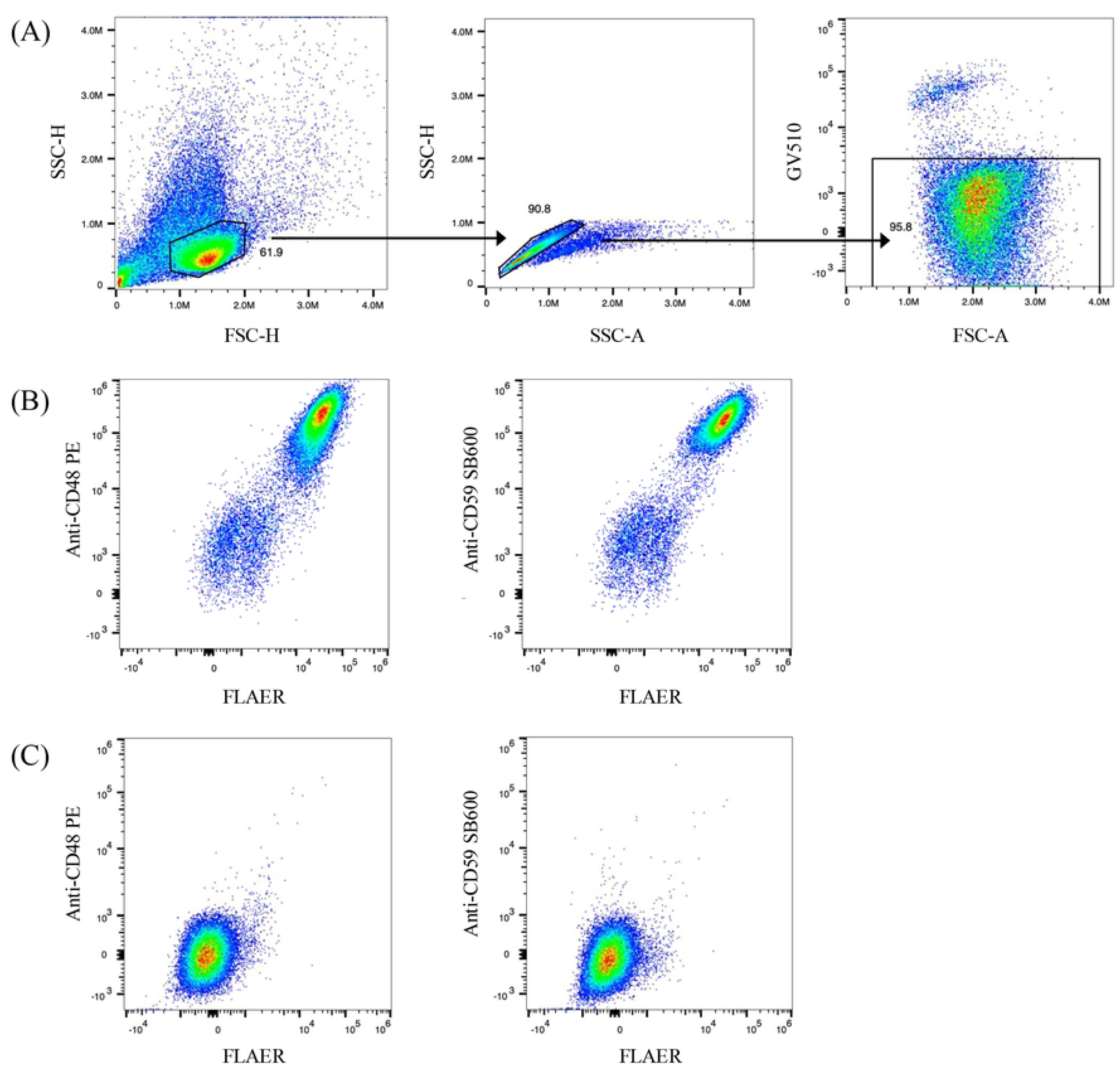
Flow cytometry of GPI, CD48, and CD59 in parental Jurkat and S12. (A) Gating strategy used to identify live, single cells of the parental Jurkat cell line. Gates first identify the cells of interest followed by excluding doublets and dead cells. (B) Plots of the singlet, live gated Jurkat population. (C) Plots of the singlet, live gated S12 population.

### Loss of GPI-AP in S12 is not caused by mutation in PIGA gDNA or mRNA exons

To examine the cause of GPI-anchor deficiency in the S12 cell line, molecular analysis of *PIGA* exons 2-6 using genomic DNA was performed. Exon 1 is small and non-coding and was not examined. No variants were detected in a comparative analysis of S12 *PIGA* gDNA amplicons to the NCBI reference sequence (NG_009786.1) using a long-read Oxford Nanopore Technologies (ONT) workflow (S1, S2, S3 and S4 FASTA files) No variants were detected in an analogous analysis of Jurkat *PIGA* gDNA (S5, S6, S7 and S8 FASTA files). However, a single nucleotide polymorphism was found between gDNA exons 2 and 3 (intron 2) in both S12 and parental Jurkat (S1 and S5 FASTA files). Additionally, no variants were detected in a comparative analysis of a *PIGA* mRNA amplicon spanning the middle of exon 2 through exon 6 to the NCBI reference sequence (NM_002641.3) in either S12 or Jurkat via the ONT workflow (S9 and S10 FASTA files).

### S12 displays significantly higher expression of CD3 and TCRαβ than parental Jurkat

Flow cytometry revealed that S12 had increased levels of both CD3 and TCRαβ relative to parental Jurkat (Fig 3). Quantitative analysis of the MFIs demonstrated a 47% increase in TCRαβ expression and a 29% increase in CD3 expression in S12 compared to parental Jurkat. The differences in CD3 and TCRαβ expression were significant between the two cell lines (p = 0.004 and 7.089 x 10^-7^, respectively). Samples were stained in triplicate, and the experiment was repeated (see file S11 MFI for data).

**Fig 3.**
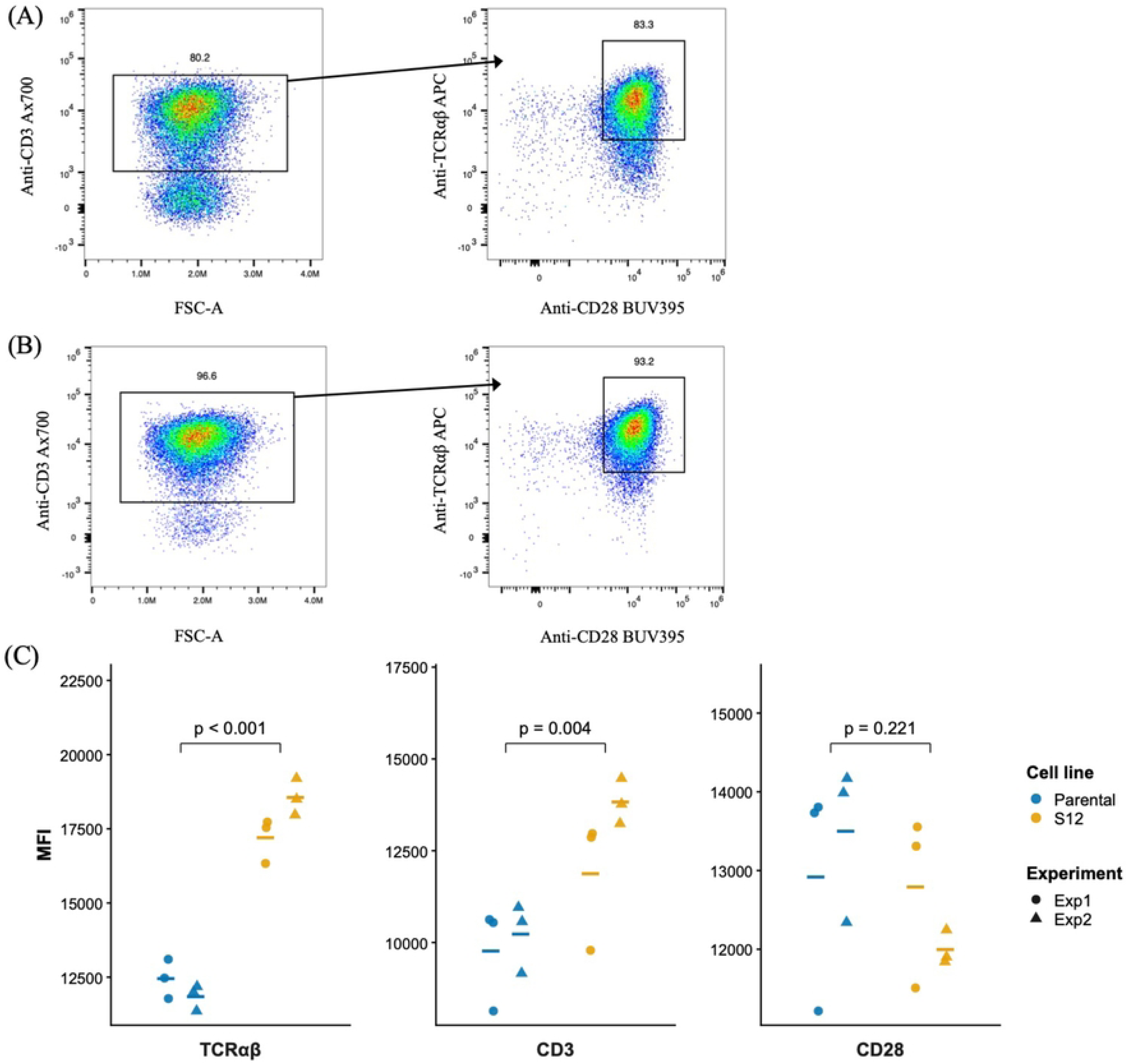
Flow cytometric analysis of CD3, CD28, and TCRαβ expression in parental Jurkat and S12. (A) The left plot displays live, parental Jurkat singlets gated as previously described (Fig. 2). The right plot shows the CD3+ parent Jurkat cells. CD3, CD28, and TCRαβ positive gates were determined using fluorescent minus one (FMO) controls. (B) The left plot displays live, S12 singlets gated as previously described. The right plot shows the CD3+ S12 cells. The gates were determined using FMO controls as previously described. (C) Individual MFI of CD28, CD3, and TCRαβ surface expression of each cell line measured in triplicate in two independent experiments are shown. The mean of the experiments is shown as a dash.

### No significant difference in IL-2 production between S12 and parental Jurkat

IL-2 production was measured by ELISA in supernatants collected 24 hours following stimulation with anti-CD3/CD28 beads or PMA/ionomycin in parental Jurkat, S12, and human PBMCs. Stimulation with the beads did not induce detectable IL-2 production in either parental Jurkat or S12. Under the same conditions, PBMCs produced substantial levels of IL-2. In contrast, PMA/Ionomycin stimulation induced IL-2 production in both Jurkat and S12. Across both statistical approaches, the overall contrast comparing parental Jurkat to S12, averaged across all three stimulation conditions, was not statistically significant (Fig. 4) (sqrt-LMM: estimate = 4.6, p = 0.360; asinh-LMM: estimate = +0.17, p = 0.267) (see file S12 IL2 ELISA for data).

**Fig 4.**
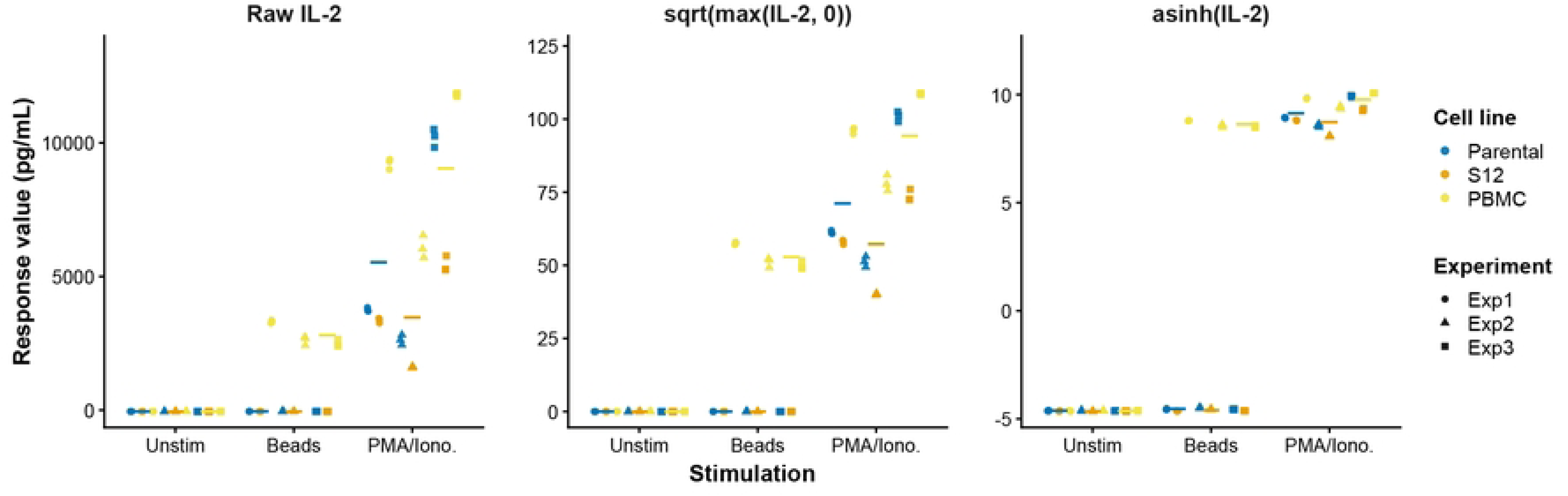
IL-2 production of parental Jurkat and S12. Raw concentrations (Raw IL-2), square-root transformation (sqrt(max(IL-2, 0))), and inverse hyperbolic sine transformation (asinh(IL-2)) of IL-2 in supernatants of Jurkat, S12 or PBMC cultured with anti-CD3/anti-CD28 Dynabeads (1:1 ratio), PMA + Ionomycin, or unstimulated detected by human IL-2 ELISA. The dash represents the mean of the respective cell line and condition.

## Discussion

This study examined the impact of GPI-deficiency on CD3, CD28, and TCRαβ expression and found that GPI-deficient T cells have significantly higher expression of CD3 and TCRαβ when compared to GPI-producing controls. In addition, IL-2 production by GPI-deficient T cells remains intact following PMA/ionomycin stimulation compared to GPI-producing controls.

Given the previously established role of lipid rafts in organizing TCR signaling complexes, GPI-deficiency would be expected to impair downstream TCR signaling (10, 11, 13). Our S12 cell line retained the ability to produce similar IL-2 levels to the parental Jurkat cell line following stimulation with PMA and ionomycin. PMA and ionomycin act downstream of TCR signaling at PKCθ and calcineurin, respectively (19). Upon TCR signaling, PKCθ localizes to membrane lipid rafts which are dependent on GPI (20). This may explain why S12 cells have slightly lower IL-2 production compared to the parental Jurkat cell line. This difference in IL-2 production would be expected to increase upon signaling that originates more upstream as it would involve more lipid raft-dependent actions. As to the increased CD3 and TCRαβ expression, it has been previously shown that GPI-deficient T cells undergo delayed TCR capping and internalization (13). This delayed TCR capping and internalization could explain the significantly higher TCRαβ expression of S12 cells compared to parental Jurkat cells. CD3 capping may be affected similarly.

Prior studies have shown that GPI-deficient T cells can both enhance and impair TCR signaling and subsequent IL-2 production. As previously mentioned, engagement of GPI-APs in a Jurkat cell line has been shown to enhance TCR signaling and IL-2 production (12). Consistent with this finding, GPI-deficient T cells from patients with PNH exhibited impaired responses to TCR signaling (13). Inconsistent with this finding, GPI-deficient murine T cells exhibited higher production of IL-2 upon TCR stimulation (21). Based on these findings, GPI-deficiency can have a spectrum of effects on T cell function via signaling through the TCR. This study demonstrates that GPI-deficiency has no significant effects on IL-2 production from signals originating at PKCθ and calcineurin - helping narrow down where in the TCR signaling pathway GPI-deficiency impacts T cell function.

Overall, there is very limited research on either TCR expression or functionality of GPI-deficient T cells. Most GPI-deficiency research focuses on GPI-deficient erythrocytes from patients with PNH. More studies like ours could help elucidate the mechanisms behind the paradoxical immune responses commonly seen in patients with PNH (22, 23). In addition, T cells with *in vivo* mutation of the *PIGA* gene and disrupted biosynthesis of GPI anchors are being studied in surrogate selection, a method that uses somatic mutation as a surrogate marker for *in vivo* T cell proliferation to investigate clinical disorders with immunological features (24). *PIGA* is a housekeeping gene on the X chromosome that encodes one subunit of an enzyme required for the biosynthesis of GPI anchors (25). Since *PIGA* is an X-linked gene, a single mutation may show a dominant effect throughout the cell, and a mutation causing dysfunction of *PIGA* will disrupt many surface markers on the cell. T cells with *in vivo* somatic mutations, including *in vivo PIGA* mutant T cells, are enriched in T cells undergoing *in vivo* clonal proliferation (26). The expanding T cell clones are expected to include TCR-defined PIGA mutant T cell clones with a *PIGA* mutation as well as T cells with the same TCR and no mutation (wild-type) (26). Thus, TCR analysis of *in vivo PIGA* mutant T cells may identify *in vivo* clonally amplified T cells for additional functional characterization (26). The TCR and functional data from this report will inform those studies.

There are several limitations present in this study. First, all experiments were run using immortalized cancer T cell lines which may not fully replicate primary T cells or T cells *in vivo*. Second, a single method, i.e., anti-CD3/anti-CD28 beads, was tested to stimulate the parental Jurkat and S12 cell lines through the TCR in this report. We did not explore if alternative methods for TCR engagement, e.g., soluble or plate-bound antibodies, would induce a cellular response from Jurkat or S12. It has been previously observed that crosslinking the TCR on Jurkat and major histocompatibility complex class II on antigen presenting cells with staphylococcal enterotoxins (SE) elicits a robust cytokine response from Jurkat (27–29). Thus, the response of our in-house Jurkat and S12 to SE stimulation warrants further investigation. Finally, the mutation/mechanism behind S12’s GPI-deficiency was not identified. However, the GPI-deficiency of S12 was a stable phenotype that was maintained in our lab for >5 months.

Future studies are proposed to extend these findings to primary human T cells. Findings from the Jurkat and S12 cells reported here will inform those studies. Additionally, these two cell lines could be used to investigate the unique selection and expansion of GPI-deficient T cells that occurs *in vivo* in patients (30).

## Conclusions

GPI-deficient Jurkat T cells have: 1) increased expression of CD3 and TCRαβ; 2) similar levels of CD28 expression; and 3) intact IL-2 production following PMA/ionomycin stimulation compared to a GPI-producing parental Jurkat.

## Acknowledgements

The authors thank the University of Wisconsin Carbone Cancer Center (P30 CA014520) and the William S. Middleton Memorial Veterans Hospital, Madison, WI. The authors also thank the UWCCC Flow Cytometry Laboratory (instrument grant #1S10OD018202-01), the UWCCC Cancer Pharmacology Laboratory, and the UWCCC Translational Research Initiatives in Pathology (TRIP) Laboratory, supported by the UW Department of Pathology and Laboratory Medicine, UWCCC and the Office of The Director - NIH (S10 OD023526) for its Cell Line Authentication Service, for use of facilities and services. Oxford Nanopore sequencing was performed at the University of Wisconsin-Madison Biotechnology Center DNA Sequencing Core Facility (RRID:SCR_017759). Fig. 1 was created in BioRender, Glass, W. (2026) https://BioRender.com/kuehg4a.

## Supporting information

**S1 Fig. Single nucleotide polymorphism within intron 2 of *PIGA* in S12 and Jurkat.** The exon 2 sequence is marked with a solid underline and the ATG start codon is marked with a wave underline and highlighted in blue. In the sequence represented here, bases 5’ (upstream) of exon 2 originate from intron 1, whereas bases 3’ (downstream) of exon 2 originate from intron 2. An adenine base change in S12 and Jurkat compared to a cytosine base in the Reference Sequence is denoted in lowercase and highlighted in yellow. This single base change, at position 1211, is within intron 2 (between exons 2 and 3) of the *PIGA* sequence. The values in the right margin denote the number of bases specific to the amplicons represented in this figure, not to the *PIGA* gene itself. Sequences were aligned with Clustal Omega (15) and an asterisk (*) indicates 100% base conservation.

**S1 FASTA File. S12 gDNA exon 2 FASTA file**

**S2 FASTA File. S12 gDNA exon 3 FASTA file**

**S3 FASTA File. S12 gDNA exons 4 and 5 FASTA file**

**S4 FASTA File. S12 gDNA exon 6 FASTA file**

**S5 FASTA File. Jurkat gDNA exon 2 FASTA file**

**S6 FASTA File. Jurkat gDNA exon 3 FASTA file**

**S7 FASTA File. Jurkat gDNA exons 4 and 5 FASTA file**

**S8 FASTA File. Jurkat gDNA exon 6 FASTA file**

**S9 FASTA File. S12 mRNA mid exon 2 through exon 6 FASTA file**

**S10 FASTA File. Jurkat mRNA mid exon 2 through exon 6 FASTA file**

**S11 MFI data file. Flow cytometry Median Fluorescence Intensity values of TCR**αβ**, CD28 and CD3 surface protein expression by S12 and Jurkat cell lines.**

**S12 IL2 ELISA data file. IL2 secreted by S12, Jurkat or human PBMC after stimulation with anti-CD3/CD28 beads PMA/Ionomycin or unstimulated.**

